# Meta-Analysis of Wild Relatives and Domesticated Species of Rice, Tomato, and Soybean Using Publicly Available Transcriptome Data

**DOI:** 10.1101/2025.06.01.656526

**Authors:** Makoto Yumiya, Hidemasa Bono

## Abstract

The domesticated species currently available in the market have been developed through the breeding of wild relatives. Breeding strategies using wild relatives with high genetic diversity are attracting attention as an important approach for ad-dressing climate change and ensuring sustainable food supply. However, studies examining gene expression variation in multiple wild and domesticated species are few. Therefore, we aimed to investigate the changes in gene expression associated with domestication. We performed a meta-analysis of public gene expression data of domesticated species of rice, tomato, and soybean and their presumed ancestral species. In wild relatives, the expression of genes involved in osmotic, drought, and wound stress tolerance was upregulated. In domesticated species, upregulated expression was observed in genes related to auxin and those involved in the efflux of heavy metals and harmful substances. These findings provide insights into how domestication influences changes in crop traits. Thus, our findings may contribute to rapid breeding and development of new varieties capable of growing in harsh natural environments. Hence, a new cultivation method called “de novo domestication” has been proposed, which combines the genetic diversity of currently unused wild relatives and wild relatives with genome editing technologies that enable rapid breeding.

## 1. Introduction

Owing to global warming, extreme weather events, and changes in the Earth environment, the yields of major global crops such as maize, rice, and soybeans could de-crease 12–20% by the end of this century [1–4]. Additionally, with an increase in global population, food demand is expected to rise significantly, and by 2050, global food demand is projected to be approximately 1.5 times higher than that in 2010 [5]. To address these issues, the development of crop varieties with desirable traits in a short period is highly anticipated. In recent years, genome editing technologies, particularly CRISPR/Cas9, have garnered significant attention. Genome editing is a technology that enables the targeting of specific genes or nucleotide sequences to induce loss-of-function or introduce genes derived from other organisms. Using this genome editing technology, new varieties of various organisms, such as tomatoes with high gamma-aminobutyric acid content and red sea bream (Madai) with increased edible parts, have been developed [6,7]. Additionally, “*de novo* domestication” efforts utilizing wild relatives, which are genetic resources that have been underutilized until now, are also gaining attention for achieving sustainable crop production. The reason for this interest is because the currently distributed domesticated species have been selectively bred with a particular focus on yield, resulting in concerns about their low genetic diversity [8–15]. To implement such methods effectively, obtaining insights into the differences in traits between domesticated species and their wild counterparts is essential. However, studies examining gene expression variation in multiple wild and domesticated species are few.

Meta-analysis is a valuable method that integrates multiple research findings to provide new insights. Meta-analyses utilizing public gene expression data have been performed and reported in literature [16]. Additionally, the number of gene expression datasets registered in public databases is expected to increase in the future [17]. Therefore, the reliability of the analysis results is expected to increase as the number of datasets available for the meta-analysis increases. Given this background, we performed a meta-analysis using gene expression data from wild and domesticated species of rice, tomato, and soybean registered by multiple research groups in public databases to identify the gene groups specifically expressed in wild relatives and crops.

This study aimed to investigate the changes in gene expression associated with domestication and provide insights for developing new crop improvement strategies. Although the species analyzed in this study were not closely related and the number of gene ex-pression datasets used was limited, this analysis provides valuable insights by focusing on changes in gene expression between wild and domesticated species during domestication.

## 2. Materials and Methods

### 2.1 Curation of Public Gene Expression Data

RNA sequencing (RNA-seq) data were obtained from public databases, primarily the National Center for Biotechnology Information Gene Expression Omnibus (NCBI GEO) [18]. To supplement the datasets not available in NCBI GEO, additional RNA-seq data were collected from published studies available online. In NCBI GEO, we performed searches using the scientific names of wild relatives, specifically “*Oryza rufipogon*” and “*Glycine soja.*” For tomato, two wild relatives, “*Solanum pennelli*i” and “*Solanum arcanum,*” were utilized. The domesticated species paired with wild relatives and used in this analysis were “*Oryza sativa japonica*” for rice, “*Solanum lycopersicum*” for tomato, and “*Glycine max*” for soybean. To further refine the search results, a filter for “expression profiling by high-throughput sequencing” was applied. RNA-seq data were searched using the methods described above, and datasets containing paired wild and domesticated species from the same project were curated and used for subsequent analyses. The rationale for using data from the same project was to standardize the cultivation environments and experimental conditions as much as possible, thereby reducing batch effects.

### 2.2 Gene Expression Quantification

Each RNA-seq dataset was obtained using the prefetch (version 3.0.10) and Fasterq-dump (version 3.0.10) commands from the SRA Toolkit [19]. Quality control of the raw reads and removal of adapter sequences were performed using fastp (version 0.23.4) [20]. Subsequently, the output files from the quality check were aggregated into a single file using MultiQC (version 1.18) [21]. Transcripts were quantified using Salmon (version 1.10.1) [22]. For transcript quantification, reference cDNAs obtained from Ensembl Plant were used: *Oryza sativa japonica* and *Oryza rufipogon* with IRGSP-1.0; *Solanum pennellii, Solanum arcanum*, and *Solanum lycopersicum* with SL3.0; and *Glycine soja* and *Glycine max* with *Glycine_max*_v2.1. As a result, the quantified RNA-seq data were expressed as transcripts per million (TPM). Subsequently, transcript-level TPM values were summarized at the gene level using the tximport (version 1.28.0) [23] package in R.

### 2.3 Calculation of DW-Ratio

Gene expression data were normalized to the DW-Ratio. ’D’ and ’W’ represent ’Domesticated’ and ’Wild,’ respectively. The DW-Ratio was calculated using the following equation:

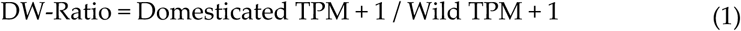

When calculating the DW-Ratio, to avoid division by zero and prevent errors caused by genes with zero expression, 1 was added to the TPM values of both domesticated species and wild relatives.

### 2.4 Classification of Differentially Expressed Genes (DEGs) Based on DW-Ratio

To evaluate genes showing expression changes between domesticated species and their wild relatives, all genes were classified into three groups. Specifically, the three groups were upregulated, unchanged, and downregulated. These groupings were determined according to preestablished thresholds. Genes were classified as upregulated if their DW-Ratio exceeded an upper threshold, downregulated if their DW-Ratio fell below a lower threshold, and unchanged if they did not meet either of these criteria. For the upregulated category, 20 thresholds ranging from 1.5-fold to 200-fold were tested, and a 2-fold threshold was adopted. For the downregulated category, 20 thresholds ranging from 1/1.5 to 1/200 were tested, and a threshold of 1/2 was adopted [24].

### 2.5 Calculation of DW-Score

To evaluate DEGs by integrating different experiments based on the DW-Ratio, a DW-Score was calculated. The DW-Score was calculated by subtracting the number of pairs classified as upregulated from the number of pairs classified as downregulated. A pair refers to a set comprising one wild relative and one domesticated species from the same project.

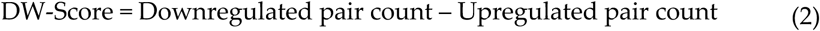

DW-Ratio and DW-Score were calculated using code from a previous study [25].

### 2.6 Gene Set Enrichment Analysis

Gene set enrichment analysis was performed using ShinyGO 0.81 [26] on the top- and bottom-ranking genes based on their DW-Scores. For the rice analysis, “*Oryza sativa japonica* Group gene IRGSP-1.0” was selected as the species, and “Gene Ontology (GO) Biological Process” was chosen as the pathway database. Default settings were used for all the other parameters. No enriched terms were observed when using species-specific annotations for tomato and soybean. Therefore, the gene IDs for both species were converted to *Arabidopsis thaliana* gene IDs, and enrichment analysis was performed using “*Arabidopsis thaliana* genes (TAIR10).”

### 2.7 Commonly Upregulated Genes in Wild and Domesticated Species

To perform cross-species analysis, gene IDs from each species were converted to their corresponding *Arabidopsis thaliana* gene IDs (TAIR10). For this process, we used Ensembl Plant BioMart [27] to create a correspondence table linking the gene IDs of rice, tomato, and soybean with the gene IDs of *Arabidopsis thaliana* (TAIR10). Using the DW-Score for each species, we performed comparisons focusing on three ranges: the top 1%, 3%, and 5% of the upregulated and downregulated genes.

## 3. Results

### 3.1 Overview of the Study

In this study, a meta-analysis was performed to identify DEGs using domesticated species and their ancestral wild relatives. Figure 1 presents an overview of the study.

**Figure 1.**
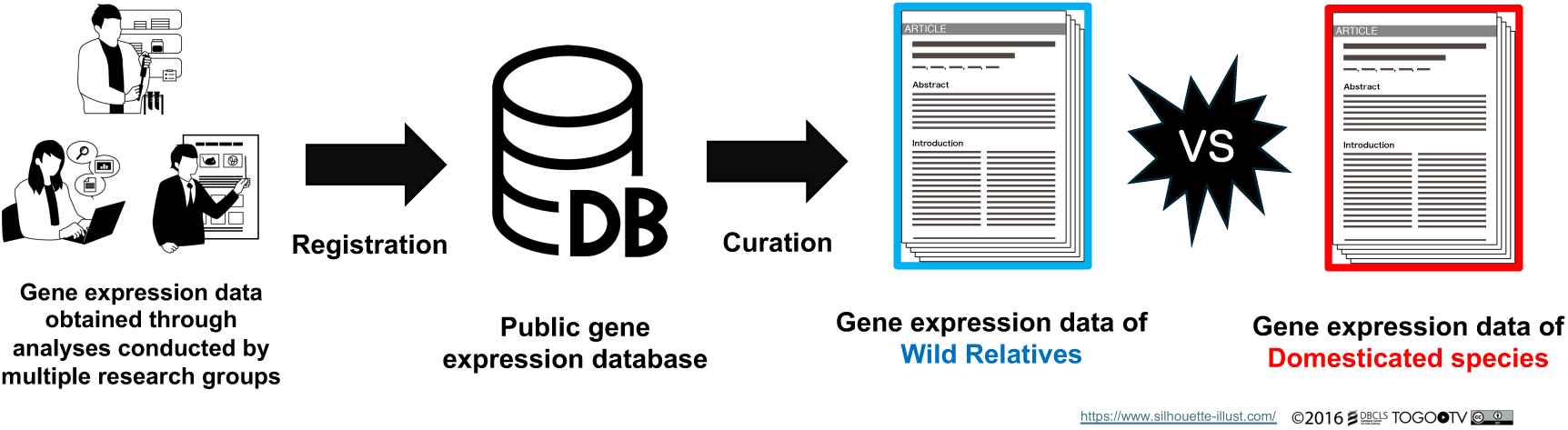
Gene expression data of wild relatives and domesticated species of rice, tomatoes, and soybeans were collected from public databases and manually curated. To ensure comparability, data only from wild and domesticated species included in the same research project were selected and paired. Subsequently, gene expression levels were quantified and differentially expressed genes between wild relatives and domesticated species were identified.

### 3.2 Curation of RNA-seq Data from Public Databases and Literature

The RNA-seq data of wild and domesticated species used in this study were primarily collected from NCBI GEO [18]. Compared to domesticated species, data on wild relatives were substantially limited. Therefore, we decided to use three species—rice, tomato, and soybean—for the analysis because data from multiple projects were available for these species. The number of samples and their tissue types used in this analysis are shown in Figure 2.

**Figure 2.**
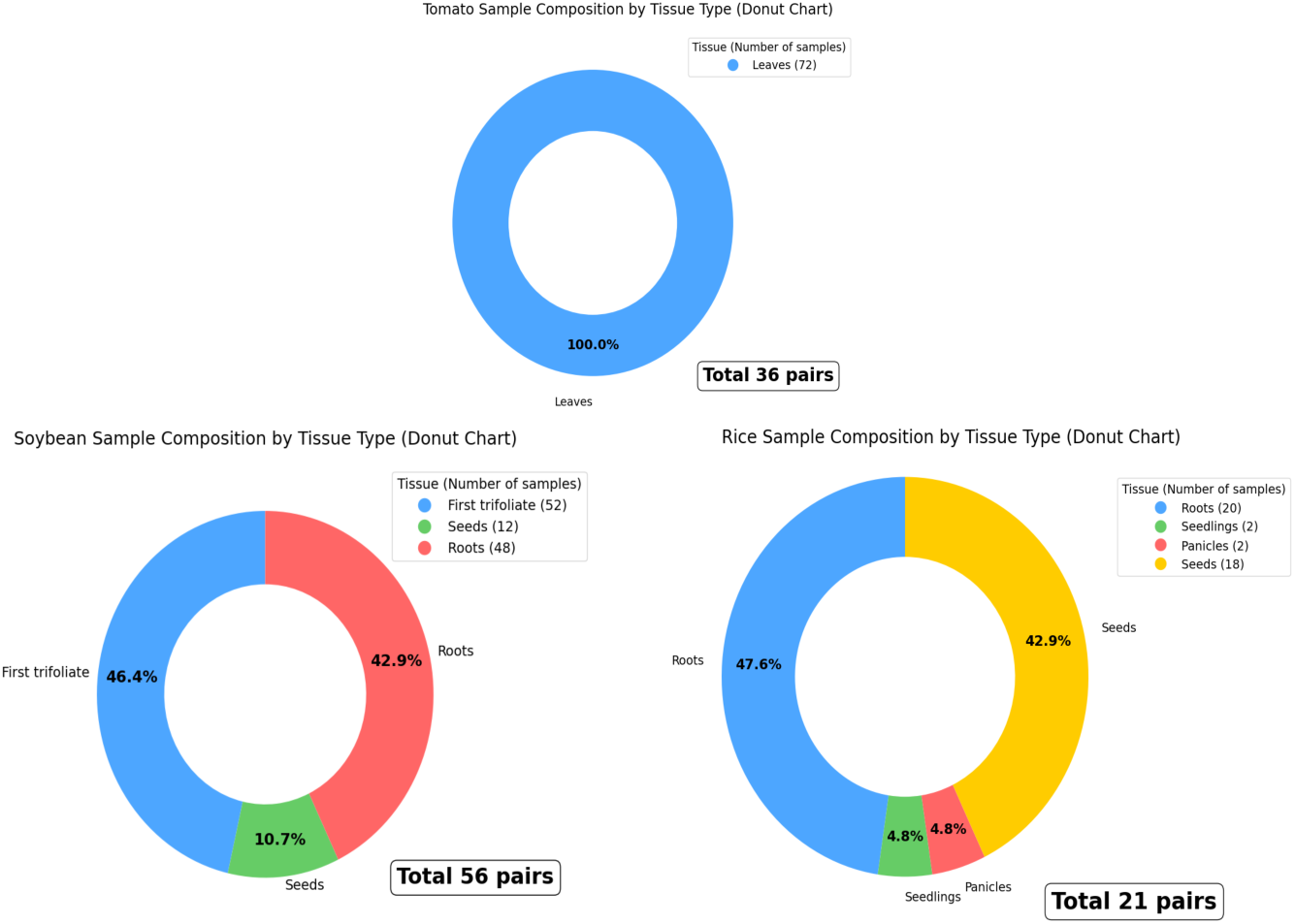
Number of plant samples and their tissue types used in this analysis. All tomato plant tissue samples were leaves, totaling 36 pairs. For soybean, the tissues included first trifoliates, seeds, and roots, with a total of 56 pairs. For rice, the tissues consisted of roots, seedlings, panicles, and seeds, totaling 21 pairs. Each pair comprised one sample from a wild relative and one sample from a domesticated species. Each sample was paired such that the wild and domesticated species shared the same tissue type and experimental conditions.

To collect RNA-seq data, keyword searches were performed using the names of three wild relatives: *Oryza rufipogon*, *Solanum pennellii / Solanum arcanum*, and *Glycine soja*. RNA-seq data were obtained from five BioProjects in the NCBI GEO and used for analysis. However, because the number of available datasets from NCBI GEO was very limited, RNA-seq data were also collected from ArrayExpress in the European Bioinformatics Institute BioStudies [28] and relevant literature. Ultimately, RNA-seq data from four BioProjects not registered in the NCBI GEO were obtained from the Sequence Read Archive [29], resulting in a total of nine BioProjects used for analysis. The metadata for the samples used in this analysis are provided in Supplementary Tables S1, S2, and S3, which correspond to *Oryza rufipogon* and *Oryza sativa japonica* (S1); *Solanum pennellii*, *Solanum arcanum*, and *Solanum lycopersicum* (S2); and *Glycine soja* and *Glycine max* (S3), respectively.

### 3.3 Classification of DEGs and Enrichment Analysis in Oryza rufipogon and Oryza sativa japonica

The expression quantification data for each species are provided in Supplementary Tables S4, S5, and S6, which correspond to *Oryza rufipogon* and *Oryza sativa japonica* (S4); *Solanum pennellii*, *Solanum arcanum*, and *Solanum lycopersicum* (S5); and *Glycine soja* and *Glycine max* (S6), respectively. Subsequently, both the expression ratio (DW-Ratio) and DW-Score of the wild and domesticated species were calculated. Based on the DW-Score ranking, lists of DEGs were obtained for both the wild and domesticated species. A threshold of two-fold and one-half-fold change was selected as the criterion for identifying genes with differential expressions. Lists of genes with DW-Scores above the two-fold and one-half-fold thresholds used in this analysis are provided in Supplementary Tables S7, S8, and S9, which correspond to *Oryza rufipogon* and *Oryza sativa japonica* (S7); S*olanum pennellii*, *Solanum arcanum*, and *Solanum lycopersicum* (S8); and Glycine soja and *Glycine max* (S9), respectively.

To investigate functional biases, we performed enrichment analysis on DEGs from pairs of *Oryza rufipogon* and *Oryza sativa japonica*, utilizing approximately the top 3% ranked using DW-Score (Figure 3a). As a supplement, for rice, an enrichment analysis was initially performed using the top approximately 1% of the DEGs based on DW-Score; however, no enriched terms were identified. Therefore, for the rice analysis only, the scope was expanded to include DEGs in approximately the top 3% based on DW-Score. In the enrichment analysis targeting genes with upregulated expression in *Oryza rufipogon*, GO terms related to environmental responses, such as “Cellular response to cold” and “response to wounding,” were found to be enriched (Figure 3b). In the results for *Oryza sativa japonica*, the GO terms related to photosynthesis and photosystems were enriched (Figure 3c). The genes included in the GO terms identified in *Oryza rufipogon* and *Oryza sativa japonic*a are listed in Supplementary Tables S10 and S11, corresponding to *Oryza rufipogon* (S10) and O*ryza sativa japonica* (S11), respectively.

**Figure 3.**
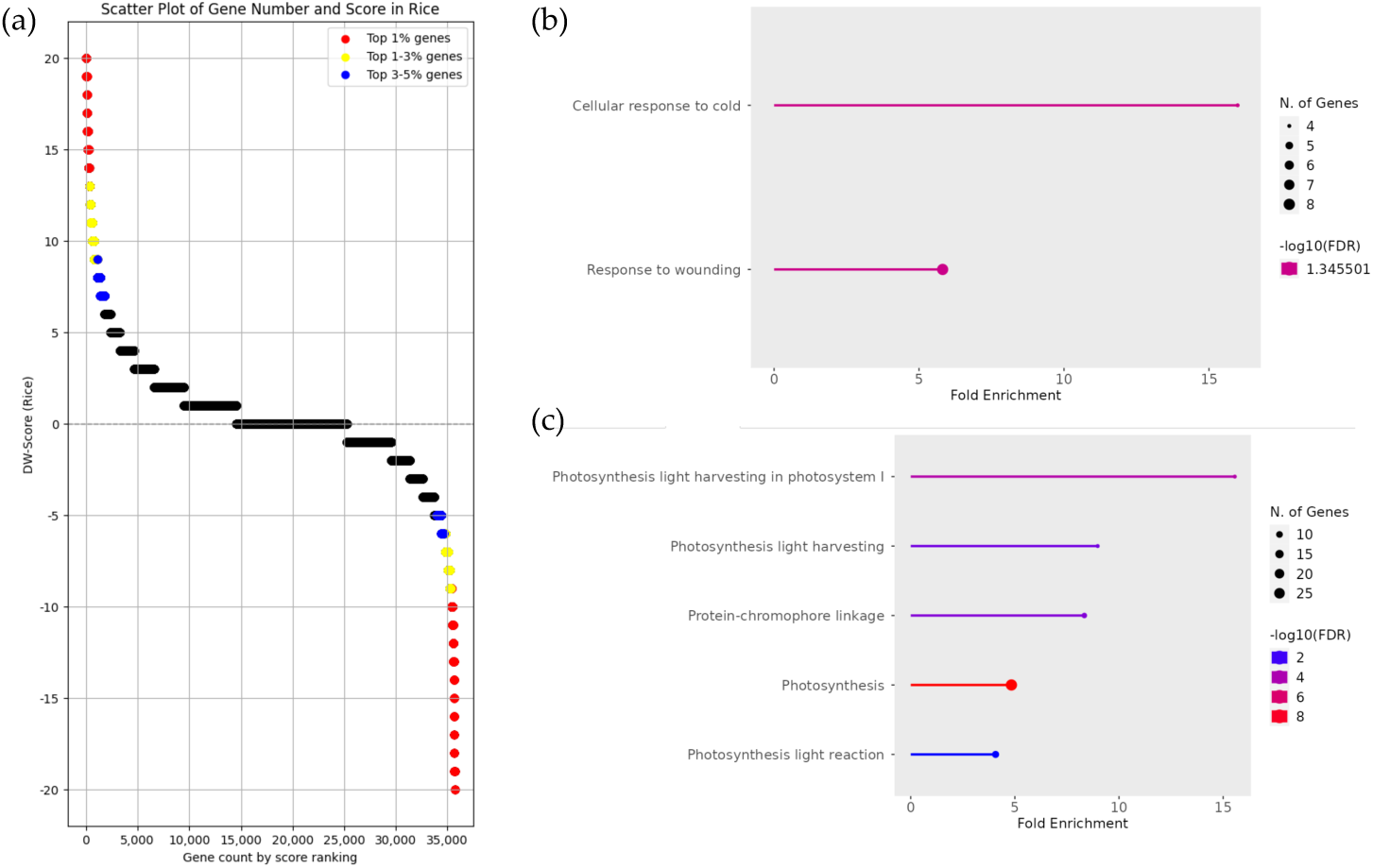
Enrichment analysis and DW-Score scatter plot of gene groups with upregulated expression in *Oryza sativa japonica* and *Oryza rufipogon*. (**a**) The scatter plot represents the DW-Score of each gene, where positive and negative scores indicate upregulated expression in *Oryza sativa japonica* and *Oryza rufipogon*, respectively. The red, yellow, and blue dots represent the top 1%, 1–3%, and 3–5% of scores, respectively. (**b**) Enrichment analysis results of gene groups with upregulated expression in *Oryza rufipogon*. The analysis was based on DW-Score, utilizing approximately the top 3% of genes, which included 898 genes (−20 ≦ DW-Score ≦ −7). (**c**) Enrichment analysis results of gene groups with upregulated expression in *Oryza sativa japonica*. The analysis was based on DW-Score, utilizing approximately the top 3% of genes, which included 1,073 genes (9 ≦ DW-Score ≦ 20).

### 3.4 Classification of DEGs and Enrichment Analysis in Solanum pennellii / Solanum arcanum and Solanum lycopersicum

In the enrichment analysis for tomato, as mentioned in the Materials and Methods section, tomato gene IDs were converted to *Arabidopsis thaliana* gene IDs, and the enrichment analysis was performed accordingly.

Enrichment analysis was performed on DEGs from pairs of *Solanum pennellii*, *Solanum arcanum*, and *Solanum lycopersicum*, utilizing approximately the top 1% ranked using DW-Score (Figure 4a). In the enrichment analysis of gene groups with upregulated expression in *Solanum pennellii* and *Solanum arcanum*, GO terms such as “ascorbate glutathione cycle” and “purine nucleoside transmembrane transport” were found to be enriched. Additionally, GO terms related to stress responses, such as “response to hydrogen peroxide” and “response to reactive oxygen species,” were also enriched (Figure 4b). In *Solanum lycopersicum*, GO terms related to sulfur metabolism and plant hormones, such as “Sulfate reduction,” “Jasmonic acid metabolic process,” and “Salicylic acid metabolic process,” were enriched (Figure 4c). The genes included in the GO terms identified in *Solanum pennellii*, *Solanum arcanum*, and *Solanum lycopersicum* are listed in Supplementary Tables S12 and S13, corresponding to *Solanum pennellii and Solanum arcanum* (S12) and *Solanum lycopersicum* (S13), respectively.

**Figure 4.**
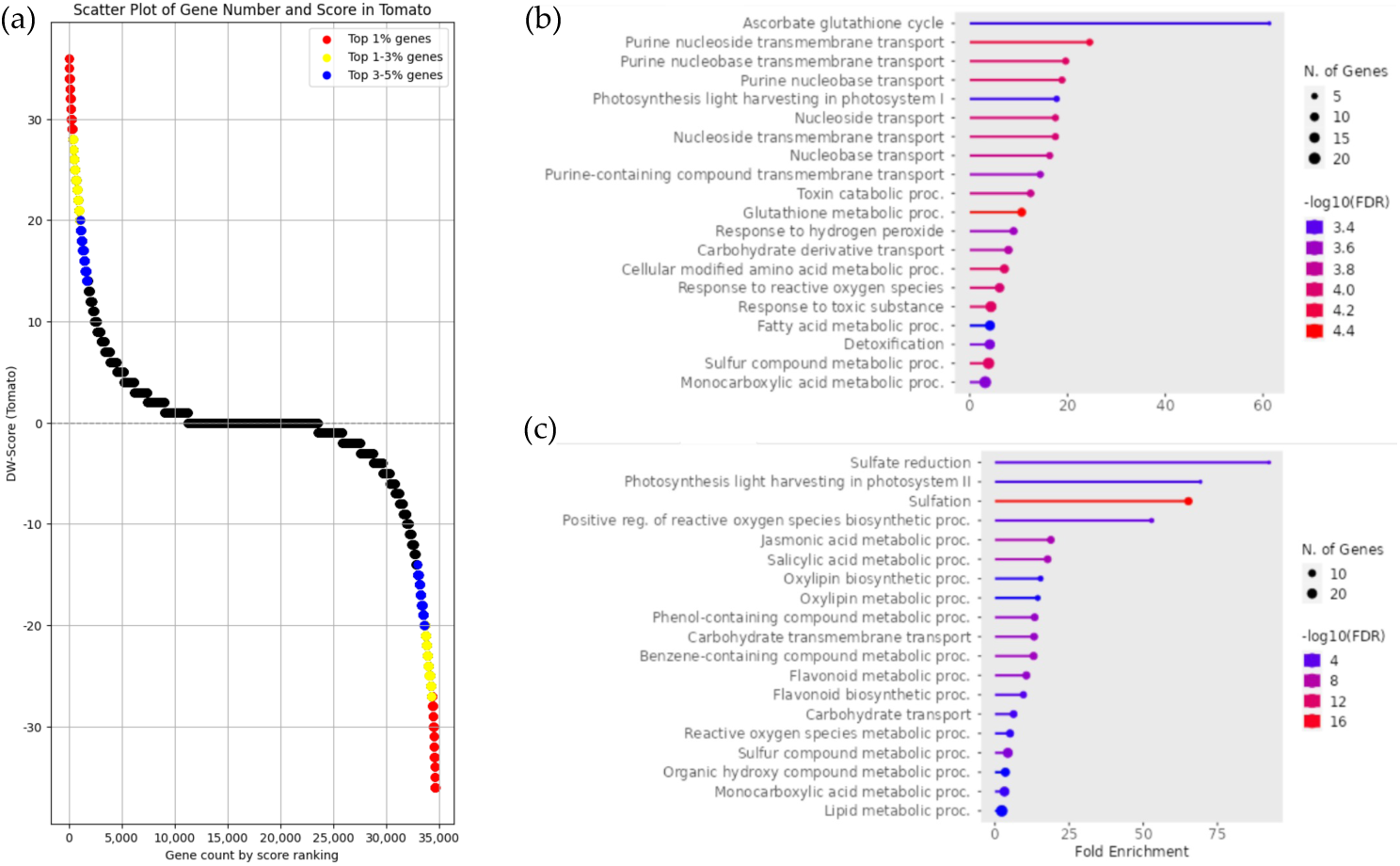
Enrichment analysis and DW-Score scatter plot of gene groups with upregulated expression in *Solanum lycopersicum* and *Solanum pennellii / Solanum arcanum*. (**a**) The scatter plot represents the DW-Score of each gene, where positive and negative scores indicate upregulated expression in *Solanum lycopersicum* and *Solanum pennellii / Solanum arcanum*, respectively. The red, yellow, and blue dots represent the top 1%, 1–3%, and 3–5% of scores, respectively. (**b**) Enrichment analysis results of gene groups with upregulated expression in *Solanum pennellii* / *Solanum arcanum*. The analysis was based on DW-Score, utilizing approximately the top 1% of genes, which included 391 genes (−36 ≦ DW-Score ≦ −27). (**c**) Enrichment analysis results of gene groups with upregulated expression in *Solanum lycopersicum*. The analysis was based on DW-Score, utilizing approximately the top 1% of genes, which included 382 genes (28 ≦ DW-Score ≦ 36).

### 3.5 Classification of DEGs and Enrichment Analysis in Glycine soja and Glycine max

Similar to tomato, soybean gene IDs were converted to *Arabidopsis thaliana* gene IDs, and enrichment analysis was performed.

Enrichment analysis was performed on DEGs from pairs of *Glycine soja* and *Glycine max* utilizing approximately the top 1% ranked using DW-Score (Figure 5a). In *Glycine soja*, similar to the results observed in other wild relatives, GO terms such as “Response to oxidative stress” and “Cellular detoxification,” which contribute to defense mechanisms against environmental stress, were enriched. Additionally, a larger number of genes were classified under these terms (Figure 5b). In *Glycine max*, the enrichment analysis revealed that GO terms associated with secondary metabolites, such as triterpenoid and isoprenoid biosynthesis, as well as pathways related to energy metabolism, were enriched (Figure 5c). The genes included in the GO terms identified in *Glycine soja* and *Glycine max* are listed in Supplementary Tables S14 and S15, corresponding to *Glycine soja* (S14) and *Glycine max* (S15), respectively.

**Figure 5.**
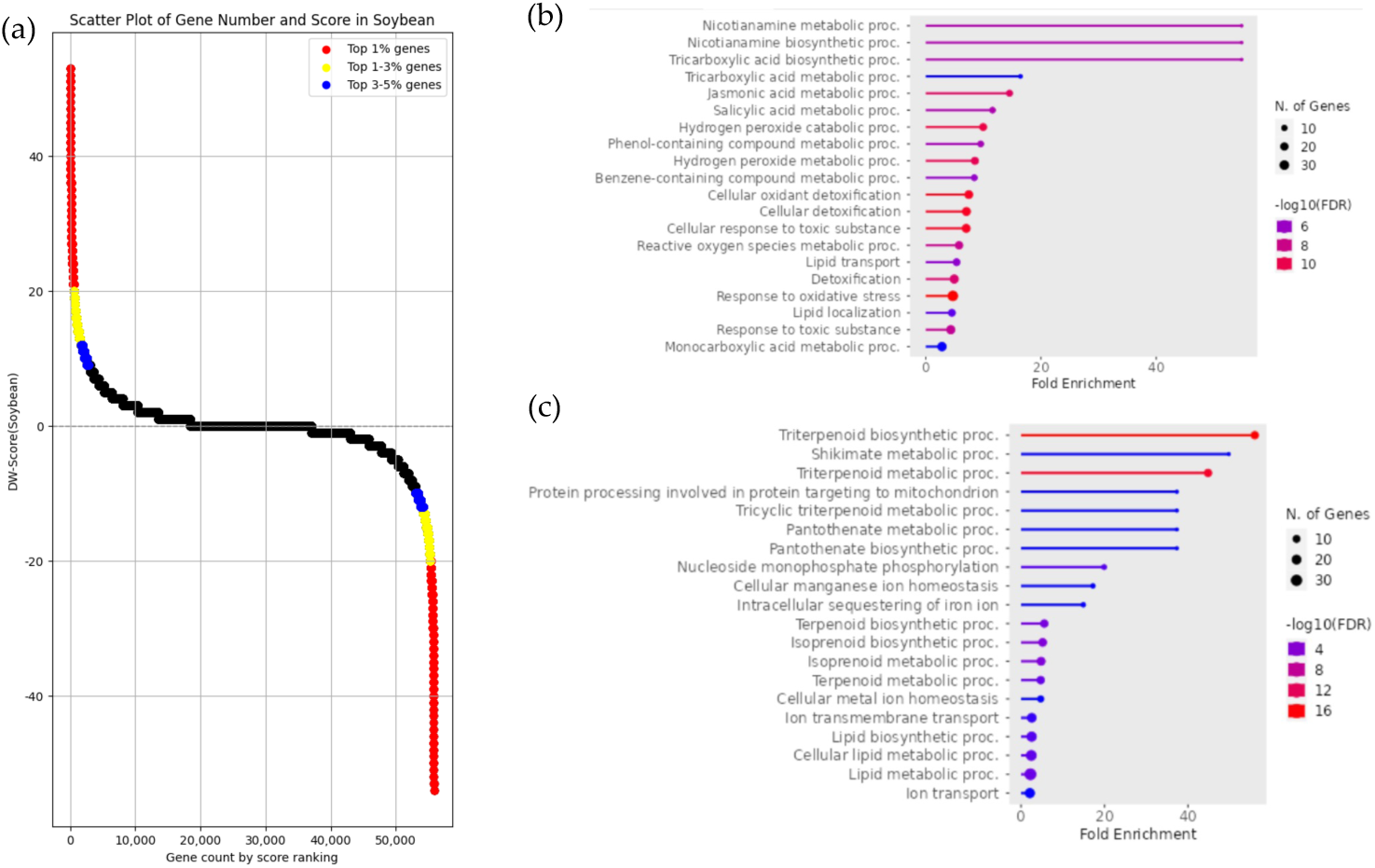
Enrichment analysis and DW-Score scatter plot of gene groups with upregulated expression in *Glycine max* and *Glycine soja*. (**a**) The scatter plot represents the DW-Score of each gene, where positive and negative scores indicate upregulated expression in *Glycine max* and *Glycine soja*, respectively. The red, yellow, and blue dots represent the top 1%, 1–3%, and 3–5% of scores, respectively. (**b**) Enrichment analysis results of gene groups with upregulated expression in *Glycine soja*. The analysis was based on DW-Score, utilizing approximately the top 1% of genes, which included 582 genes (−54 ≦ DW-Score ≦ −20). (**c**) Enrichment analysis results of gene groups with upregulated expression in *Glycine max*. The analysis was based on DW-Score, utilizing approximately the top 1% of genes, which included 590 genes (20 ≦ DW-Score ≦ 53).

### 3.6 Common DEGs in Wild and Domesticated species

To investigate genes commonly differentially expressed between wild and domesticated species of rice, tomato, and soybean, the gene IDs of each species were converted to *Arabidopsis thaliana* gene IDs. The correspondence tables for each gene ID created using BioMart [27] from Ensembl Plants focusing on the top 5% of genes by score for each species (Figure 3a, 4a, 5a) are provided in Supplementary Tables S16, S17, S18, S19, S20, and S21, corresponding to *Oryza rufipogon* (S16), *Oryza sativa japonica* (S17), *Solanum pennellii / Solanum arcanum* (S18), *Solanum lycopersicum* (S19), *Glycine soja* (S20), and *Glycine max* (S21), respectively.

Eighteen genes were commonly upregulated in approximately the top 5% of DW-Scores for wild relatives, while 36 genes were identified for domesticated species (Figure 6a, b). These genes are listed in Table 1 and 2. Enrichment analysis results for the genes that were commonly upregulated in wild relatives and domesticated species are shown in Figure 6c, d. In wild relatives, GO terms related to environmental stress responses were enriched, consistent with the previous enrichment analysis results. However, in domesticated species, GO terms related to chemical compound export and detoxification, as well as the regulation of hormone responses, were enriched, differing from the enrichment analysis results observed in each individual domesticated species.

**Figure 6.**
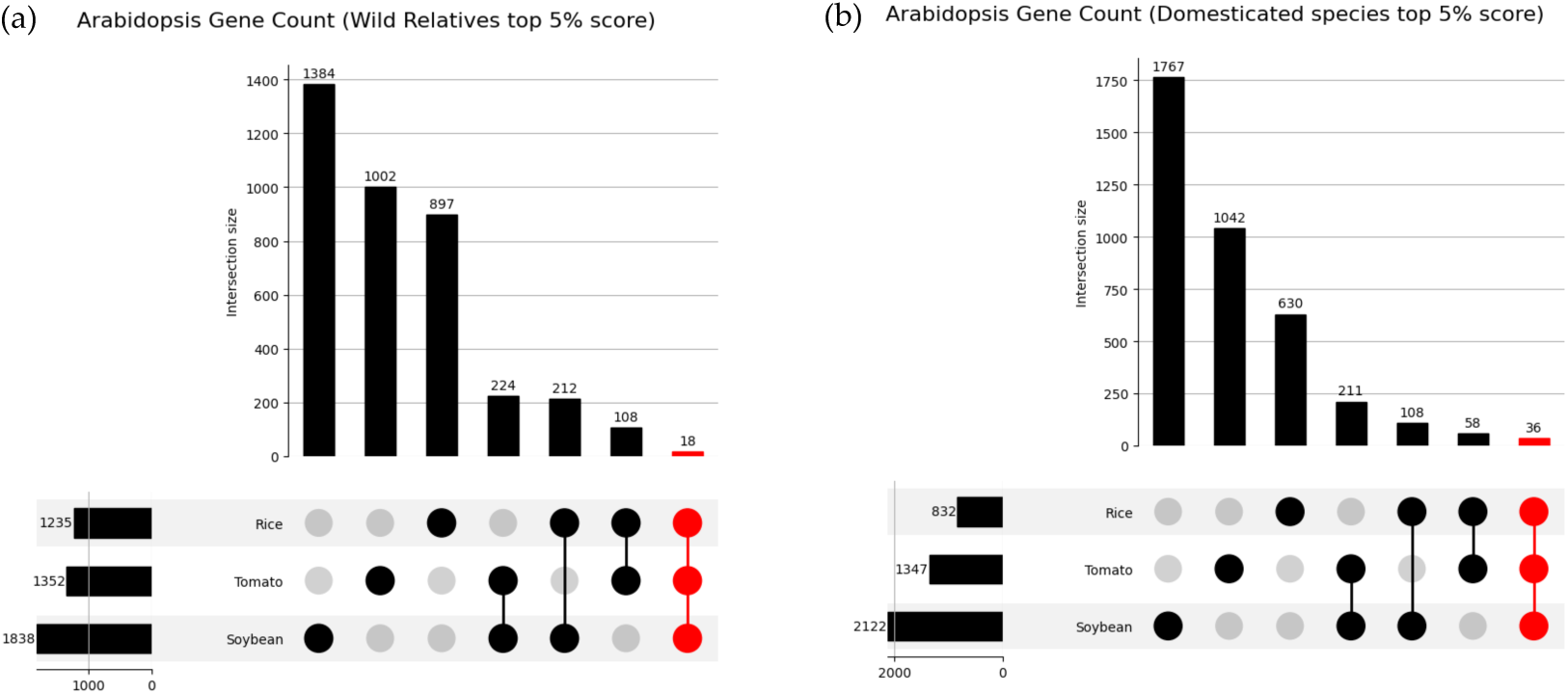
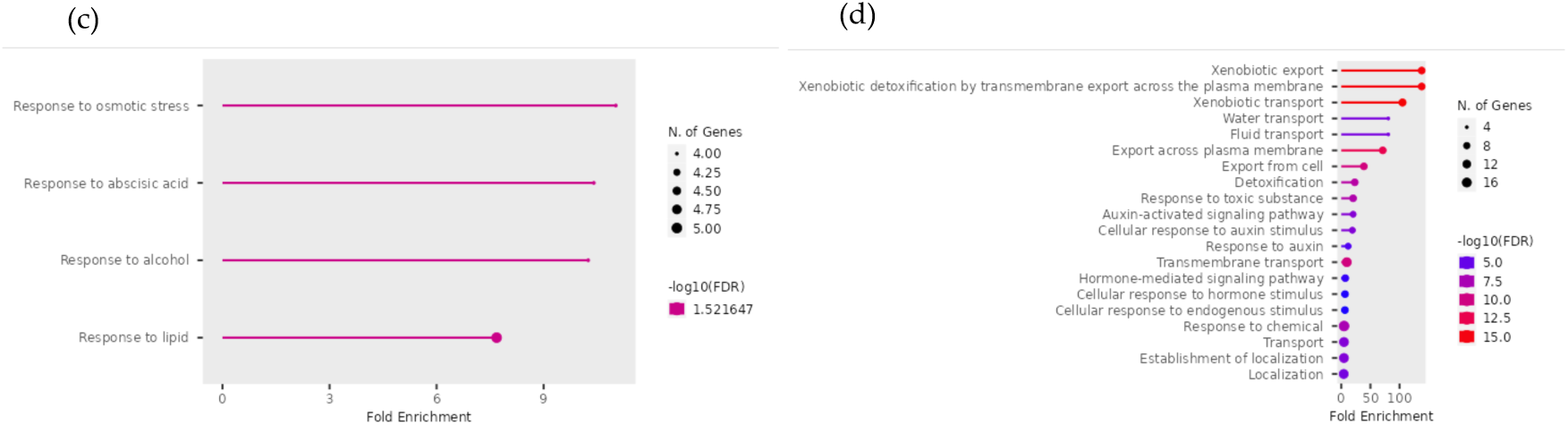
(**a**) UpSet plot of top 5% genes in wild relatives of rice, tomato, and soybean. (**b**) UpSet plot of top 5% genes in domesticated species of rice, tomato, and soybean. Enrichment analysis was performed on the top 5% of genes in both wild and domesticated species. (**c**) Enrichment analysis results for the 18 genes commonly upregulated in wild relatives. (**d**) Enrichment analysis results for the 36 genes commonly upregulated in domesticated species.

**Table 1.**
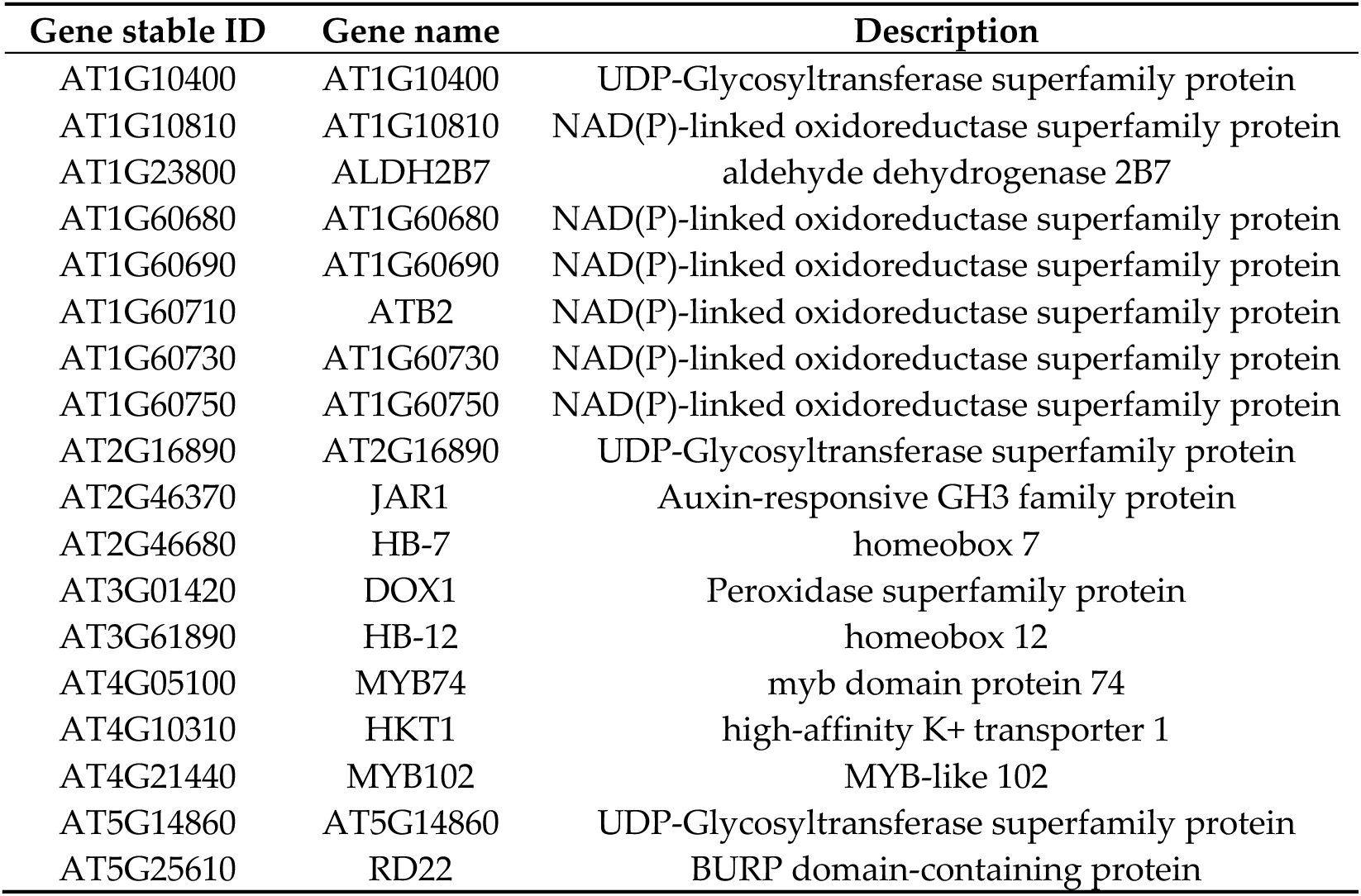
Genes commonly upregulated in wild relatives (*Oryza rufipogon*, *Solanum pennellii, Solanum arcanum*, and *Glycine soja*).

**Table 2.**
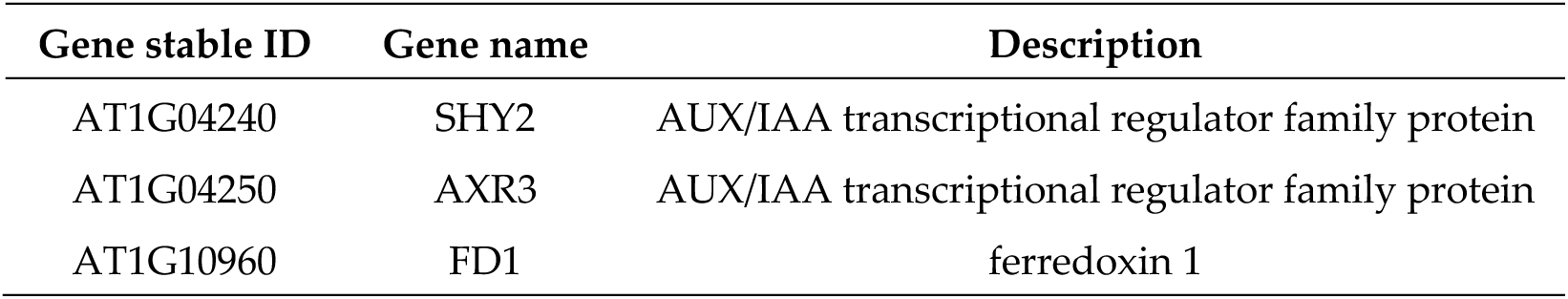
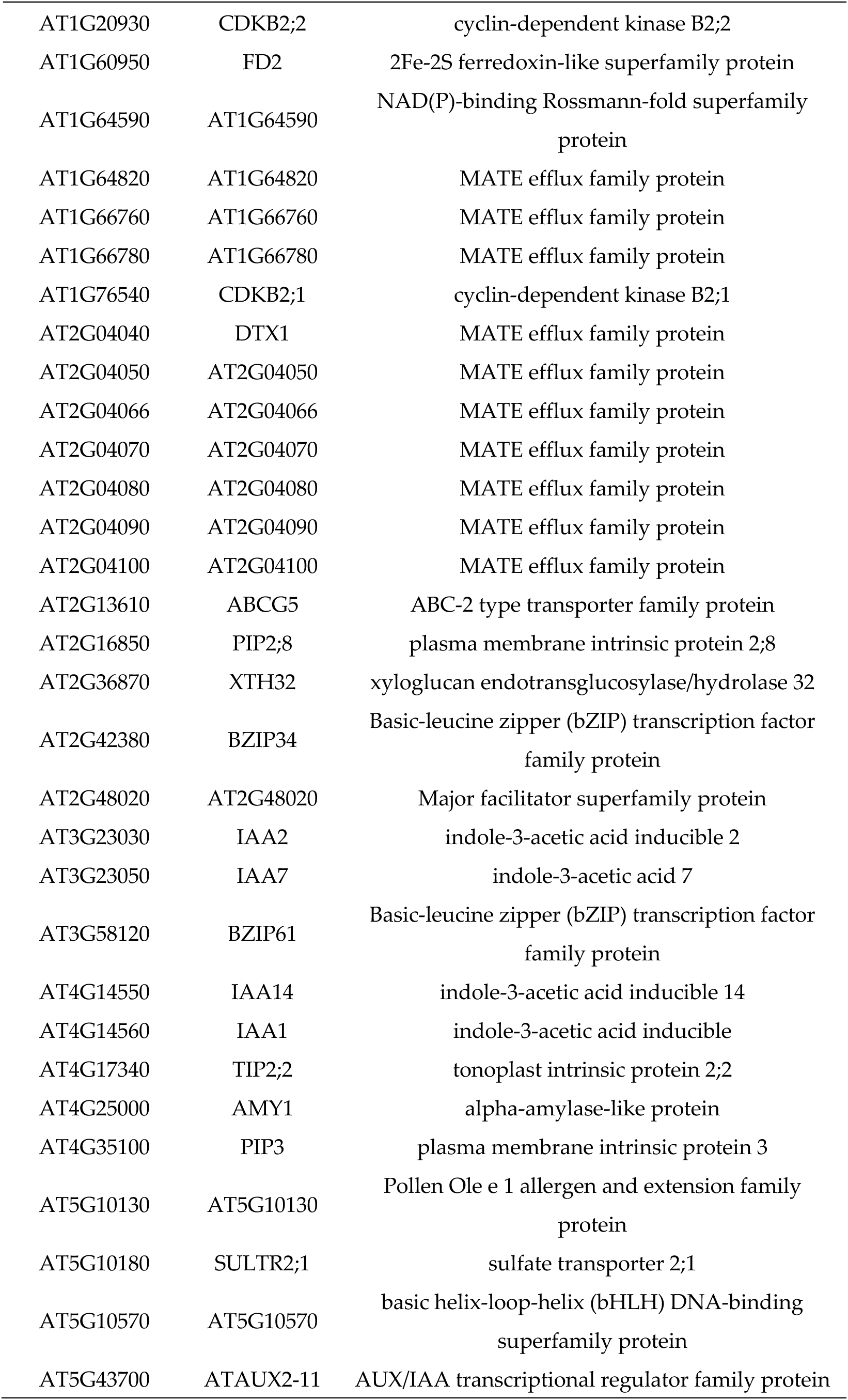
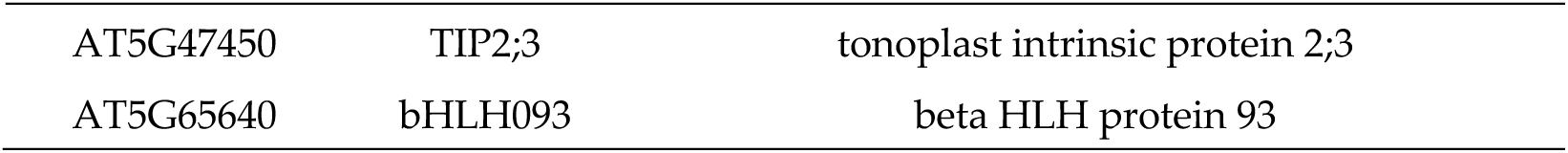
Genes commonly upregulated in domesticated species (*Oryza sativa japonica*, *Solanum lycopersicum*, and *Glycine max*).

## 4. Discussion

In this study, we obtained gene expression data for wild and domesticated species of rice, tomato, and soybean from public databases and investigated the DEGs resulting from domestication effects. For this purpose, we utilized the DW-Ratio and DW-Score as analytical metrics to perform a meta-analysis comparing gene expression data from multiple research projects. Enrichment analysis was performed to investigate the characteristics of DEGs in both wild and domesticated species. Additionally, to identify genes that showed common differential expression across different species, the gene IDs of rice, tomato, and soybean were converted to *Arabidopsis thaliana* gene IDs. Based on these analyses, wild relatives included gene groups involved in environmental stress responses, enabling plants to adapt to harsh conditions. This finding supports the previously reported high environmental adaptability of wild relatives. In contrast, domesticated species contained genes involved in detoxification and export of chemical compounds. This is likely due, in large part, to the increased use of chemical fertilizers in crop cultivation. Based on these findings, the meta-analysis utilizing gene expression data from public databases suggests the high environmental adaptability of wild relatives and changes in crop traits and characteristics resulting from domestication.

### 4.1 Genes Commonly Upregulated Across Wild Relatives

In wild relatives, 18 genes were commonly upregulated within the top 5% of DW-Scores. The gene groups included in this analysis were associated with four GO terms: Response to osmotic stress (GO:0006970), Response to abscisic acid (GO:0001101), Response to alcohol (GO:0097305), and Response to lipid (GO:0033993), as shown in the enrichment analysis results in Figure 5c. A notable result is the enrichment of the abscisic acid response term, which plays a central role in responses to various stresses, such as drought, salinity, and low temperatures, across multiple species [30–33]. Additionally, GO terms related to the osmotic stress response and maintenance of cellular homeostasis were also enriched. Thus, the genes commonly upregulated in wild relatives included gene groups involved in environmental stress responses, suggesting that wild relatives possess a higher capacity to adapt to harsh natural environments than domesticated species.

Subsequently, we investigated whether the individual genes listed in Table 1—those commonly upregulated across wild relatives—have been previously reported to primarily function in environmental stress tolerance. HKT1 is a key ion transporter involved in the plant salt stress response and salt tolerance. It primarily limits the translocation of Na⁺ from roots to leaves and stems, thereby contributing to the maintenance of ion homeostasis [34]. A study comparing salt tolerance between HKT1 knockout plants and those with phloem-specific overexpression of HKT1 reported that overexpression lines exhibited reduced Na⁺ translocation to the leaves, whereas knockout lines showed significant Na⁺ accumulation in the leaves. Additionally, plants with overexpressed HKT1 produce more seeds and have higher overall yield under saline conditions compared to control plants [35]. Therefore, the molecular mechanism of salt tolerance mediated by HKT1, which exhibits upregulated expression in wild relatives, holds promise for developing crop varieties capable of thriving in saline-affected soils while maintaining superior yields.

The RD22 gene functions as a molecular link connecting abscisic acid (ABA) signaling and abiotic stress responses, and plays a critical role in plant drought stress adaptation [36]. In *Arabidopsis thaliana*, RD22 exists as a single-copy gene, whereas certain plant species, such as grapevines, possess multiple paralogs, forming an expanded RD22 family [37]. The expression of RD22 gene is regulated by two transcription factors. When plants are exposed to drought stress, ABA is synthesized in the initial phase, triggering the production of MYB2 and MYC2—the transcription factors responsible for RD22. These factors promote RD22 gene expression, resulting in the synthesis of RD22 gene products that confer drought tolerance in plants [36,38–40].

The transcription factors HB-7 and HB-12 belong to the homeodomain-leucine zipper subfamily I. HB-12 contributes to enhanced seed production under water stress conditions, while HB-7 is involved in leaf development and photosynthesis promotion in mature plants [41]. These two genes are cooperatively regulated depending on developmental stages and environmental conditions, modulating processes associated with plant growth and water stress responses. DOX1 is an enzyme that catalyzes the initial oxidation of fatty acids and possesses diverse functions, including pathogen defense, aphid-induced wound response, and protection against oxidative stress and cell death [42–44].

MYB74 and MYB102 belong to the R2R3-MYB transcription factor family and regulate stress responses and other plant-specific processes. MYB102 is involved in the osmotic stress response and wound signaling pathway [45]. Specifically, in experiments investigating the feeding effects caused by *Pieris rapae* larvae, MYB102 knockout mutants exhibited accelerated larval development rates and considerably higher pupation rates than control plants [46]. Thus, MYB102 contributes to herbivory resistance. Similar to MYB102, MYB74 is associated with environmental stress tolerance. Specifically, MYB74 overexpression enhances osmotic stress tolerance. However, MYB74 overexpression lines exhibit detrimental effects on growth compared with control plants [47]. These findings show that although MYB74 overexpression enhances stress tolerance, particularly to osmotic stress, it negatively impacts plant growth. The present analytical results, showing higher MYB74 expression in wild relatives and lower expression in domesticated species, suggest that domestication prioritizes yield. This implies a trade-off relationship; domesticated species lost the high stress tolerance inherent in wild relatives but gained increased yield, as supported by functional studies of MYB74.

In summary, the gene function analysis of wild relatives revealed that genes contributing to traits essential for survival in harsh environments, including biotic and environmental stresses, were highly expressed. Furthermore, the fact that genes highly expressed in wild relatives exhibited reduced expression in domesticated species suggests that domestication may have led to the gradual loss of these stress-resistance genes. Therefore, breeding approaches utilizing wild relatives—which retain the high stress tolerance lost in domesticated species—hold potential as valuable genetic resources for developing crop varieties adapted to increasingly harsh environmental conditions.

### 4.2 Genes Commonly Upregulated Across Domesticated Species

In domesticated species, the terms identified in the enrichment analyses of individual wild relatives were rarely observed. This difference can be attributed to the fact that rice, tomato, and soybean are not closely related species and each species has undergone domestication and breeding to acquire different characteristics and traits. Conversely, the fact that genes involved in environmental stress tolerance were commonly detected even in analyses targeting non-closely related plant species suggests that these genes may play important roles across a wide range of plant species.

Next, we investigated the functions of genes that were commonly upregulated in domesticated species. The gene list for domesticated species included several auxin-related genes, such as SHY2, IAA3, AXR3, IAA7, IAA14, IAA1, and ATAUX2-11. Auxins are plant hormones involved in various functions, including promotion of cell elongation and regulation of plant growth and development [48,49]. Although these genes have not been reported to directly contribute to increased crop yield, the observed upregulation of auxin-related genes in domesticated species probably reflects domestication-driven selection.

Among the genes showing upregulated expression in domesticated species, a particularly distinctive feature was the inclusion of multiple multidrug and toxic compound extrusion (MATE) family genes. ALF5, a member of the MATE family, confers resistance to tetramethylammonium. Furthermore, studies suggest that engineering plants to overexpress specific MATE proteins could enable their growth in chemically contaminated soils [50]. The MATE family genes that exhibited upregulated expression in this study were considerably enriched in GO terms related to chemical export and detoxification (Figure 6d), including Detoxification (GO:0098754), Response to toxic substance (GO:0009636), Xenobiotic export (GO:0046618), Xenobiotic detoxification by transmembrane export across the plasma membrane (GO:1990961), Xenobiotic transport (GO:0042908), Export from cell (GO:0140352), and Export across plasma membrane (GO:0140115). One of the genes included in these terms, DTX1, mediates the efflux of the heavy metal cadmium as well as toxic compounds [51]. One possible reason for the increased expression of MATE gene family members in domesticated species is the change in the soil environment resulting from the increased use of chemical fertilizers. The adverse effects of heavy metal and pesticide accumulation in modern agricultural soils on plant growth and health have been extensively documented in scientific studies [52] Considering these factors, agricultural environments for domesticated species, domesticated from wild ancestors, exhibit elevated heavy metal accumulation in soils compared to traditional systems, driven by increased agrochemical use (e.g., pesticides and chemical fertilizers) and pollution linked to modern agricultural practices. Consequently, we hypothesized that domesticated species exhibit an upregulated expression of genes associated with the detoxification and export of harmful substances, a phenomenon that is not observed in their wild counterparts. However, most MATE family genes, other than DTX1, remain unnamed and are likely to be understudied, warranting further detailed functional analyses.

XTH32, another gene showing upregulated expression in domesticated species, is a member of the enzyme family involved in plant cell wall remodeling. It catalyzes the cleavage and polymerization of xyloglucan, thereby contributing to plant growth and development [53]. Therefore, XTH32 probably contributes to the development of large-yielding individuals commonly observed in domesticated species. Furthermore, this gene family is involved in adaptation to external stresses and is mentioned in meta-analyses of *Arabidopsis* abiotic stress responses performed using methods similar to this analysis [54,55].

This study has several limitations that need to be taken into consideration. First, the number of datasets used for the meta-analysis was small, which may have introduced bias in the results. This is due to the current scarcity of research projects in public databases that include RNA-seq data for both wild and domesticated species. This issue will likely be resolved in the future when studies comparing wild and domesticated species are conducted and their respective RNA-seq data accumulate. Second, well-annotated transcriptome data for wild relatives were lacking. The reference transcriptome used to quantify expression levels was derived from domesticated species. Consequently, the accuracy of gene expression quantification and identification of specific genes in wild relatives may have been compromised. To address this issue, the establishment of high-quality data for wild relatives is essential. Third, when comparing genes with variable expression across different species, the analysis was standardized to *Arabidopsis* gene names. However, because tomato, rice, and soybean are not closely related, several genes could not be matched. Consequently, genes exhibiting variations in expression in both wild and domesticated species may not be reflected in the results of this study. This issue is likely to occur when standardizing gene names to those of specific species. However, improvements are expected as the identification of gene functions advances in many species beyond model organisms. Finally, because the identification of genes with different expression levels in this study did not involve statistical testing, the results should be interpreted with caution. Despite these limitations, this study is a valuable resource for understanding trait changes due to domestication and for applications in breeding, given that few studies have examined gene expression variation in multiple wild and domesticated species.

## 5. Conclusions

A meta-analysis involving multiple wild relatives and domesticated species enabled the identification of DEGs associated with the effects of domestication. In wild relatives, increased expression of multiple genes that contribute to environmental stress tolerance was observed. These findings suggest that wild relatives have the potential to grow and survive in harsh environmental conditions. This finding also supports recent trends in breeding methods that utilize valuable traits of wild relatives. In contrast, in domesticated species, particularly rice, an upregulation of photosynthesis-related genes was observed. Since photosynthesis is closely associated with crop yield, this result may reflect the selection and breeding of high-yielding individuals during the domestication process. In the three domesticated species, the genes commonly upregulated included those involved in chemical response, efflux, and detoxification. These results reflect the impact of increased agrochemical use (e.g., pesticides and chemical fertilizers) on contemporary crop cultivation. Additionally, auxin-related genes and enzymes involved in cell-wall remodeling were identified among the upregulated genes in domesticated species, suggesting their potential contribution to yield improvement through domestication-driven selection.

The findings of this study provide valuable insights into the genetic basis of crop domestication and contribute to the identification of candidate genes for breeding that are expected to be applied in future crop improvement efforts. In particular, the utilization of wild relatives to develop new crop varieties holds significant potential as a valuable approach for addressing various challenges in future crop breeding. Although the number of species and samples used in this analysis was limited, combining analyses involving a greater diversity of species with emerging scientific technologies such as genome editing is expected to enable the development of crop varieties with desirable traits in a shorter timeframe than that required for traditional methods.

**Supplementary Materials:** The following supporting information can be downloaded at: (https://doi.org/10.6084/m9.figshare.c.7801100.v1)

Table S1: Oryza rufipogon and Oryza sativa japonica sample metadata

Table S2: Solanum pennellii・Solanum arcanum and Solanum lycopersicum sample metadata

Table S3: Glycine soja and Glycine max metadata

Table S4: TPM Data of oryza rufipogon and oryza sataiva japonica

Table S5: TPM Data of Solanum pennellii・Solanum arcanum and Solanum lycopersicum

Table S6: TPM Data of Glycine soja and Glycine max

Table S7: DW-Score Oryza rufipogon and Oryza sataiva japonica

Table S8: DW-Score Solanum pennellii・Solanum arcanum and Solanum lycopersicum

Table S9: DW-Score Glycine soja and Glycine max

Table S10: Gene List Oryza sativa rufipogon Enrichment analysis

Table S11: Gene List Oryza sativa japonica Enrichment analysis

Table S12: Gene List Solanum pennellii・Solanum arcanum Enrichment analysis

Table S13: Gene List Solanum lycopersicum Enrichment analysis

Table S14: Gene List Glycine soja Enrichment analysis

Table S15: Gene List Glycine max Enrichment analysis

Table S16: Arabidopsis thaliana - Oryza rufipogon DW-Score Top5 Percent_Gene_Correspondence

Table S17: Arabidopsis thaliana - Oryza sativa japonica DW-Score Top5 Per-cent_Gene_Correspondence

Table S18: Arabidopsis thaliana - Solanum pennellii・Solanum arcanum DW-Score Top5 Percent_Gene_Co rrespondence

Table S19: Arabidopsis thaliana - Solanum lycopersicum DW-Score Top5 Percent_Gene_Correspondence

Table S20: Arabidopsis thaliana - Glycine soja DW-Score Top5 Percent_Gene_Correspondence

Table S21: Arabidopsis thaliana - Glycine max DW-Score Top5 Percent_Gene_Correspondence

## Author Contributions

Conceptualization, M.Y. and H.B.; methodology, M.Y. and H.B.; software, M.Y.; validation, M.Y. and H.B.; formal analysis, M.Y.; investigation, M.Y.; resources, H.B.; data curation, M.Y.; writing—original draft preparation, M.Y.; writing—review and editing, M.Y. and H.B.; visualization, M.Y.; supervision, H.B.; project administration, H.B.; funding acquisition, H.B

## Institutional Review Board Statement

Not applicable.

## Informed Consent Statement

Not applicable.

## Data Availability Statement

The original data presented in the study are openly available in FigShare at https://doi.org/10.6084/m9.figshare.c.7801100.v1.

## Acknowledgments

Computations were performed using the computers at the Hiroshima University Genome Editing Innovation Center.

## Abbreviations

The following abbreviations are used in this manuscript:

RNA-seq: RNA sequencing
NCBI GEO: National Center for Biotechnology Information Gene Expression Omnibus
TPM: Transcripts per million
DW: Domesticated and Wild
DEG: Differentially expressed gene
GO: Gene ontology
ABA: Abscisic acid
MATE: Multidrug and toxic compound extrusion

